# Chronically disrupted sleep induces senescence in the visceral adipose tissue of C57BL/6 mice

**DOI:** 10.1101/2023.10.17.562803

**Authors:** Daria Timonina, Genesis V Hormazabal, Indra Heckenbach, Edward Anderton, Lauren Haky, Ariel Floro, Rebeccah Riley, Ryan Kwok, Stella Breslin, Harris Ingle, Ritesh Tiwari, Olga Bielska, Morten Scheibye-Knudsen, Herbert Kasler, Judith Campisi, Marius Walter, Eric Verdin

## Abstract

The role of sleep in systemic aging remains poorly understood, despite sleep’s essential function in preserving overall health and the prevalence of reduced sleep quality in modern society. Although reduced sleep correlates with an elevated risk of age-related diseases in humans, the mechanisms underlying this are unclear. In this study, we established a link between sleep and aging by demonstrating that disrupting sleep in C57BL/6 mice drives cellular senescence in the visceral adipose tissue. Sleep disruption also led to increased oxidative stress and DNA damage, both recognized triggers for senescence induction. Cellular senescence is implicated in numerous age-related conditions which are associated with insufficient sleep, such as cardiovascular disease, type 2 diabetes, and chronic inflammation. Our findings identify an accumulation of senescent cells in the adipose tissue, which serves as a potential target through which disturbed sleep accelerates the aging process and elevates the risk of age-related diseases.

## Introduction

Sleep is a physiological state essential for maintaining life^1,2^. Despite its importance, poor sleep quality is increasingly common. 35% of adults in the United States report sleeping less than 7 hours per night, and almost half experience fatigue during the day^3^. Sleep deficiency correlates with increased risk of pathologies such as immune dysfunction and systemic inflammation, along with elevated susceptibility to age-related diseases such as type 2 diabetes, dementia, and cardiovascular disease^4–6^. Intriguingly, aging in humans is often accompanied by a decrease in sleep quality, characterized by shortened time in deep sleep and an increase in awakenings during the sleep period^7^. Given the crucial role of sleep in preserving health, and the overlap between sleep disruption and age-related pathologies, we hypothesized that disrupted sleep contributes to accelerated aging.

Accumulation of senescent cells is a hallmark of aging. Senescent cells have been strongly implicated in numerous age-related pathologies, including those associated with sleep loss in humans, such as systemic inflammation, atherosclerosis, and type 2 diabetes^8–10^. Cellular senescence is a state of cell-cycle arrest that likely contributes to age-related tissue dysfunction by secretion of a senescence-associated secretory phenotype (SASP). This dynamic array of biologically active molecules includes pro-inflammatory cytokines, chemokines, and proteases^11^. We hypothesized that sleep disruption could contribute to disease risk by driving the accumulation of senescent cells. To test this, we used mechanical stimulation to chronically disrupt the sleep of C57BL6/J mice and systematically analyzed senescence across various tissues. Our findings revealed a robust increase in cellular senescence in the visceral adipose tissue, accompanied by DNA damage and oxidative stress, both of which can induce cellular senescence^12,13^. This study lays the groundwork for understanding how reduced sleep quality may increase the risk of age-related diseases through cellular senescence and underscores the necessity for further investigation into how sleep influences the aging process.

## Results & Discussion

### Chronic sleep disruption model in C57BL6/J mice

To elucidate the effects of chronically reduced sleep quality in adults, we used an automated mechanical system (Pinnacle Technologies) to disrupt the sleep of middle-aged (48-week-old) C57BL6/J mice for 30 days. This system comprises a cylindrical enclosure equipped with a rotating bar traversing the cage floor (Fig. 1a). The bar’s slow, gentle contact with the mice prompts them to step over it, thus disrupting their sleep without eliciting handling stress. Such methods highly disrupt sleep while minimally reducing total quantity of sleep or affecting circadian rhythm^14,15^. In our experimental design, mice were awakened approximately three times per minute throughout the duration of the experiment (Fig. 1b). Considering that C57BL6/J mice naturally sleep in 5-minute bouts, this paradigm roughly corresponds to a human being awakened twice per hour^16^. To confirm sleep disruption, we measured circulating interleukin-6 (IL-6), a marker of sleep disruption in both humans and mice, and found that mice with disrupted sleep had higher IL-6 compared to control mice (Fig. 1c)^17,18^. Control mice slept *ad libitum* in the same type of cage directly adjacent to the sleep disruption cage.

**Figure 1:**
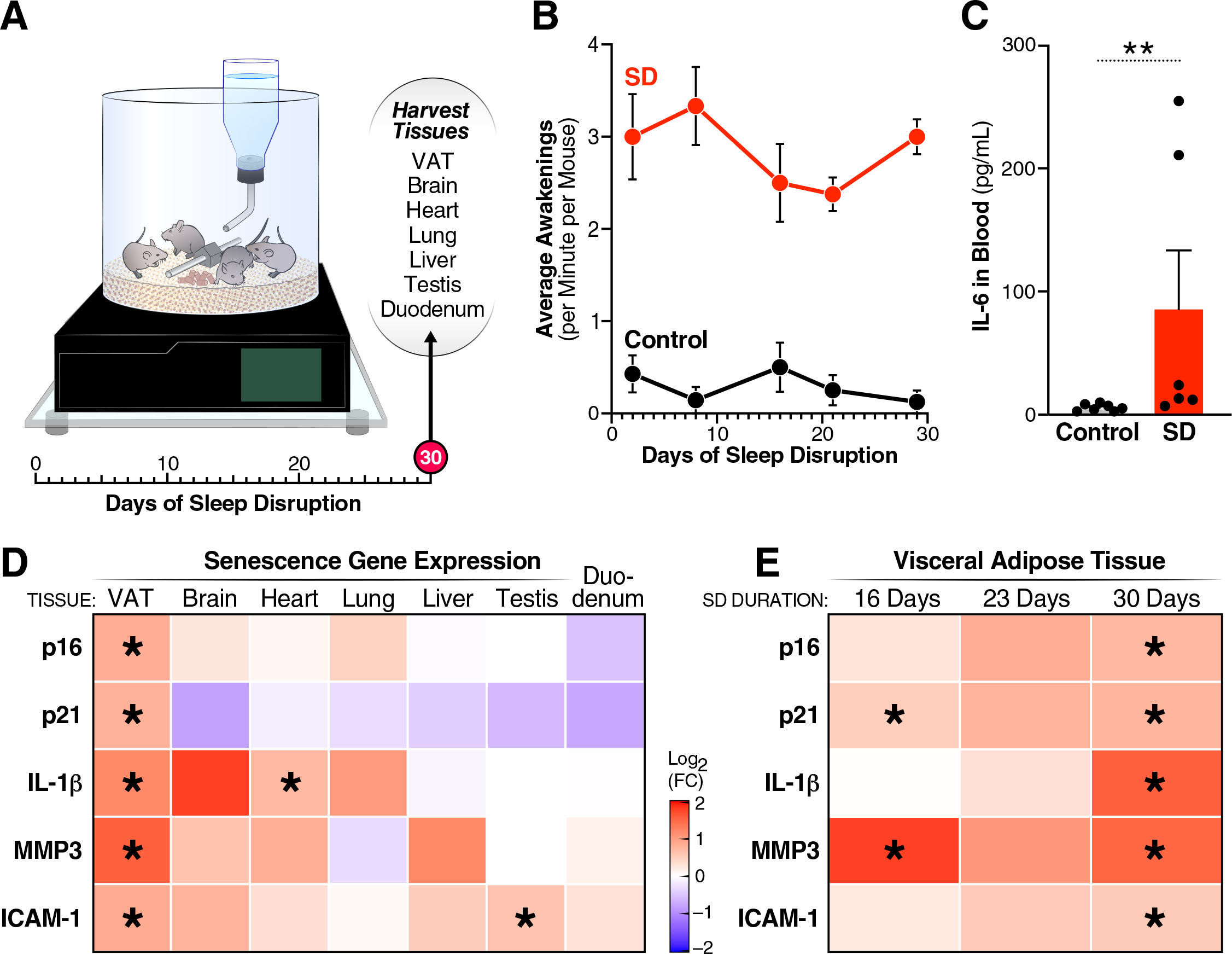
Disrupted sleep upregulates senescence gene expression. (A) Image of mechanical stimulation machine with schematic of experiment. (B) Average number of awakenings per mouse throughout course of experiment. (C) Blood level of IL-6 at SD day 30 in 48-week-old, male, C57BL/6J mice with n=7. Mean with SEM show, p value calculated using Mann Whitney test. (D) Heatmap of senescence marker/SASP factor mRNA fold change relative to controls as determined by RT-qPCR at SD day 30 in denoted tissues or (E) time course specifically in the VAT. Stars represent q values > 0.05 by Welch’s T test with 5% FDR correction using the two-stage set-up method.

### Chronic sleep disruption induces senescence gene expression

Tissue was harvested from sleep-disrupted (SD) and control mice at the end of the 30-day sleep disruption period and a panel of senescence associated genes were measured. We measured gene expression two cell-cycle arrest proteins that are commonly upregulated in cellular senescence: p16^INK4A^ (p16) and p21^Cip1/Waf1^ variant 2 (p21)^19^. Additionally, we assessed frequently reported SASP factors IL-1β, MMP3, and ICAM-1^19,20^. The visceral adipose tissue (VAT) had significantly elevated levels of these senescence-associated genes, but the brain, heart, lung, testis, liver, and duodenum did not (Fig. 1d).

We investigated if 30 days was the minimum duration of sleep disruption required increase senescence gene expression. We therefore repeated the experiment and measured senescence markers in the VAT at 16 and 23 days of sleep disruption. At 16 days, expression levels of p21 and MMP3 were marginally increased, but only at 30 days were all genes were significantly increased (Fig. 1e). This timeframe of senescence induction aligns with other paradigms that accelerate senescence in mice, such as doxorubicin treatment and ionizing radiation^21,22^. To understand if this effect was conserved in younger mice, we repeated the experiment with 26-week-old, C57BL6/J mice. Consistent with our findings in older mice, 30 days of sleep disruption caused an increase in p16, p21, IL-1β, MMP3, and ICAM-1 expression (Fig. 2a). We observed a heterogeneity of gene expression across individual mice, which may be due to variations in senescence response trajectories^23^.

**Figure 2:**
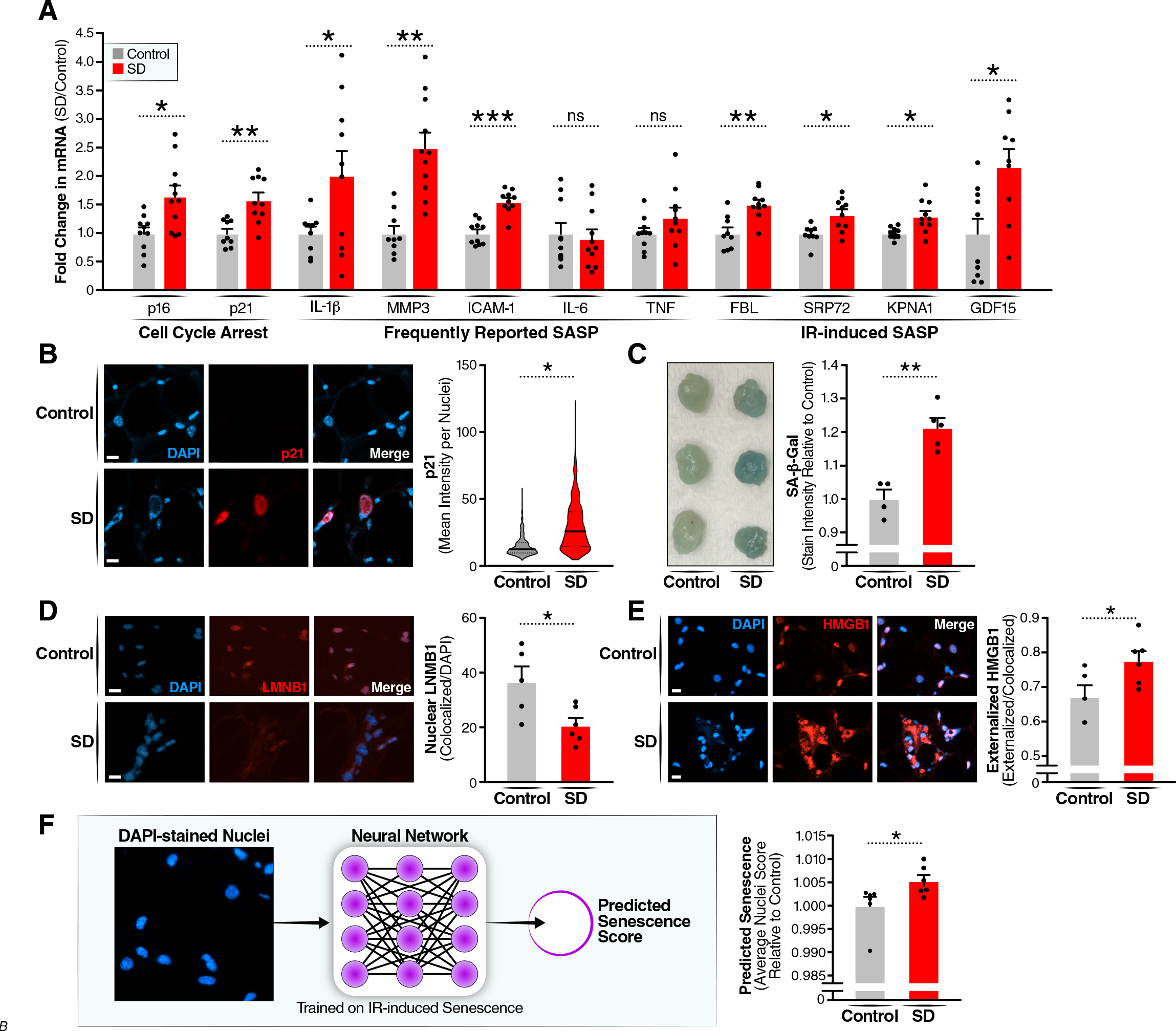
Disrupted sleep induces multiple markers of cellular senescence in the VAT. (A) RT-qPCR of senescence marker/SASP factor gene expression. Welch’s T test with 5% FDR correction, SEM. Each datapoint represents one mouse and reflects the results of two independent experiments (B) IF representative image of p21 levels in VAT with 10 μm scale bar. Graph of distribution of mean p21 intensity per nuclei. Unpaired T-test. (C) Image of VAT tissue incubated with SA-β-gal substrate and quantified for pixel intensity. (D) IF representative image of LMNB1 in VAT with 10um scale bar. Quantification of LMNB1 co-localized with DAPI normalized to DAPI area. Unpaired T-test. (E) IF representative image of HMGB1 in VAT with 10um scale bar. Quantification of HMGB1 external to Dapi normalized to HMGB1 co-localized with DAPI. Average value for each mouse plotted. Unpaired T-test with SEM. (F) Workflow diagram: DAPI images uploaded to neural network senescence (Heckenbach et al. 2022) trained of ionizing radiation (IR)-induced senescence. Neural network predicts senescence likelihood for each nuclei. Avg nuceli score for each mouse plotted. Welch’s T test.

Given the role of sleep in repairing DNA damage, we hypothesized that senescence in the VAT could be due to increased DNA damage^24^. To explore this possibility, we tested a panel of SASP factors induced by DNA-damaging irradiation^25^. These factors; FBL, SRP72, KPNA1, and GDF15, were all significantly increased in 30-day SD mice compared to control (Fig. 2a). Therefore, senescence in this model may be linked to DNA damage.

We also observed higher circulating levels of IL-6 in SD 26-week-old mice (Fig. S1). However, IL-6 mRNA expression did not increase in the VAT indicating that circulating IL-6 probably originated from elsewhere in the body (Fig. 2a). Moreover, TNFα, a cytokine secreted by certain senescent cells which increases in human blood under total sleep deprivation, remained unaltered in the VAT between control and SD mice^26^.

### Further characterization of senescence in the VAT

We next sought to determine if cellular senescence is manifested beyond changes in gene expression and to delineate the specific features of senescence induced by chronically disrupted sleep. We used immunofluorescence (IF) to detect p21 and found it significantly increased in the nucleus of SD mice VAT (Figure 2b). We also detected the following senescence markers: increased senescence-associated beta-galactosidase (SA-β-gal) activity, reduced levels of nuclear lamina protein lamin B1 (LMNB1), and relocalization of the nuclear protein HMGB1 (Fig. 2c-e). Interestingly, the increase in SA-β-gal was already detectable at day 16 and continued to rise through days 23 and 30 (Fig. S2), suggesting that SA-β-gal is an early indicator or pre-senescence marker.

To confirm our findings, we used an established and validated neural network trained to predict senescence based on nuclear morphology from irradiation-induced (IR) senescence^27^. This analysis predicted the VAT of SD mice to have more senescent cells (Fig 2f), consistent with our previous data that sleep disruption-induced senescence may stem from DNA damage.

### Oxidative stress and DNA damage are present in the VAT of SD mice

The observed similarity between the SASP profile in our model and that of irradiation-induced senescence led us to hypothesize that DNA damage could be responsible increased senescence after sleep disruption. Supporting this hypothesis, we found an increase in the DNA damage marker, phosphorylated-histone H2A.X, in the VAT of mice subjected to chronic sleep disruption (Fig 3a).

**Figure 3:**
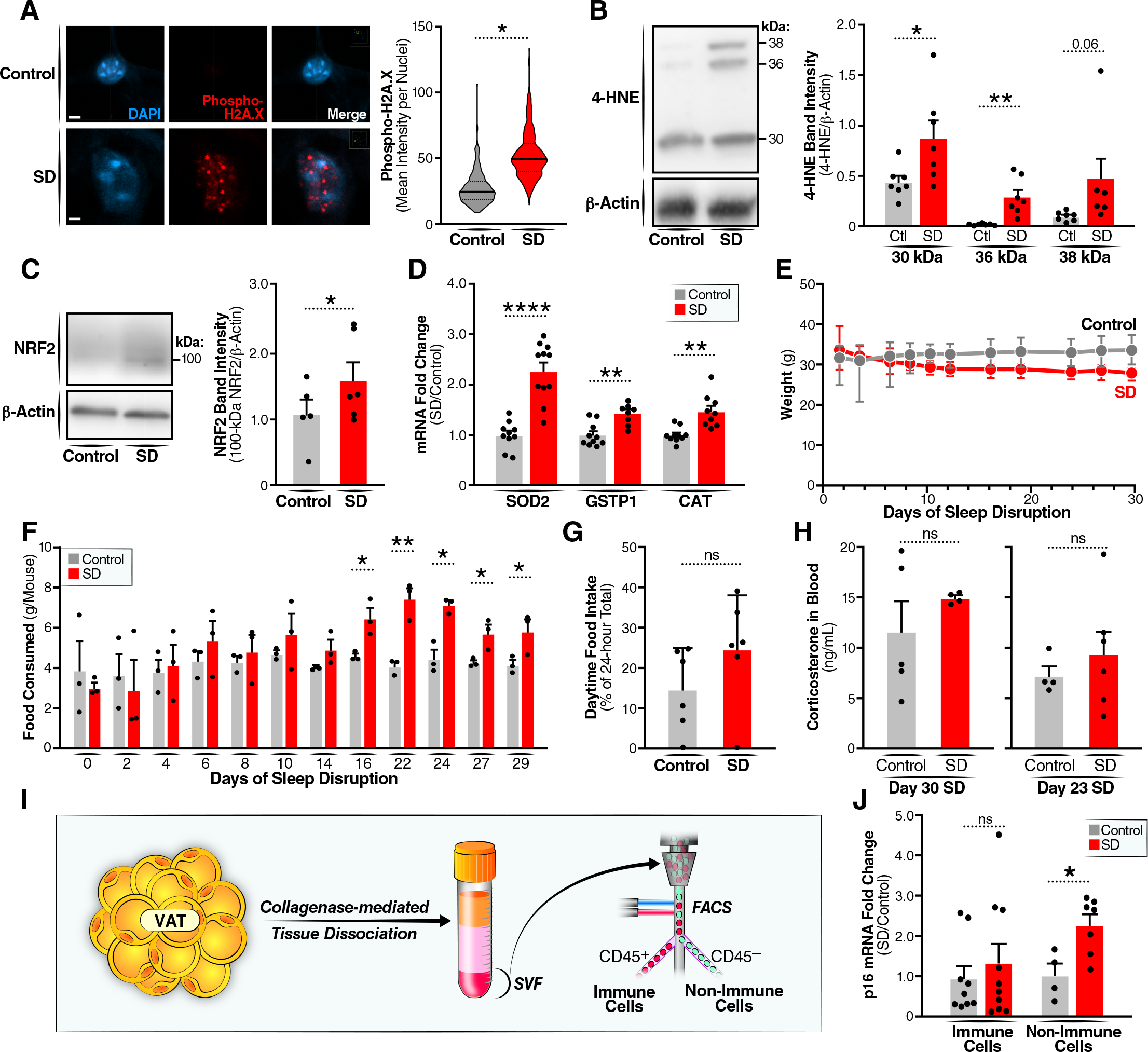
Oxidative stress and DNA damage are present in the VAT of SD mice. (A) IF representative image of phosphorylated H2A.X (Ser139/Tyr142) in the VAT with 4um scale bar. Graph of the distribution of mean intensity for γH2AX per nuclei. Unpaired T-test. (B) Western blot of 4-HNE protein levels in control or SD mouse with quantifications for each band size (C) Western blot for NRF2 protein with band quantification (D) RT-qPCR for oxidative stress response genes for each mouse. (E) Average mouse weight over the course of sleep disruption. (F) Food intake in 24hrs per mouse in control cages compared to SD cages. Measured per cage and then divide by the number of mice in that cage. (G) Amount of food eaten during lights-on (active) hours divided by food eaten in 24hrs. Measured per cage starting from SD day 16. (H) Corticosterone level in blood at 30 and 23 days into sleep disruption. (I) Diagram of workflow for sorting leukocytes from non-leukocytes (J) p16 mRNA expression in sorted cell populations. Unpaired T-test.

Sleep has been shown to mitigate oxidative stress, and therefore, we hypothesized that disrupted sleep may lead to oxidatively damaged cellular components and double-strand DNA breaks^28,29^. We explored the possibility of oxidative damage by measuring 4-hydroxynonenal (4HNE), a stable product of the oxidation of ω-6-unsaturated fatty acids, which forms HNE protein adducts and propagates oxidative stress^30^. We found increased 4HNE modifications to proteins of 30 and 36 kDa in the VAT of SD mice compared to control mice (Fig. 3b).

NRF2 protein, which accumulates in response to oxidative stress and facilitates repair of irradiation-induced DNA damage, was also elevated in VAT of SD mice (Fig 3c)^31^. Furthermore, expression of oxidative response genes superoxide dismutase type 2 (SOD2), glutathione S-transferases P1 (GSTP1), and catalase (CAT) were elevated in the VAT of SD mice compared to control (Fig. 3d). These findings indicate the presence of oxidative stress and DNA damage, which likely contribute to the initiation of senescence pathways^32^.

### Senescence is not due to weight gain

Obesity has been linked to the induction of cellular senescence^33^. However, in our study, SD mice lost an average of 4.7 grams body weight during the initial 11 days, with more substantial weight loss observed in heavier mice. After this initial period, weights remained relatively stable (Fig 3e). Prior studies on sleep disruption reported varied outcomes (e.g., weight loss, no weight change, or weight gain), depending on the experimental paradigm^34–36^. In our specific model, weight gain was not a contributing factor to the increase in cellular senescence.

Food consumption was similar between control and SD mice for the first 10 days, after which SD mice began consuming more food than control mice, a trend that became statistically significant at day 16 and continued for the remainder of the experiment (Fig 3f). Notably, the proportion of food consumed during either dark or light hours remained unchanged (Fig 3g), suggesting that increased food intake was not associated with atypical eating during regular sleeping hours. The increase in eating behavior followed by stabilization of weight likely indicates that a point of energy homeostasis was reached.

### Senescence is not due to stress or from p16-expressing immune cells

We questioned whether senescence was attributable to stress from the sleep disruption paradigm we measured corticosterone levels in the blood of mice, which are increased in mice during chronic stress. SD mice showed no increase in corticosterone levels at days 23 or 30 (Fig. 3h).

To eliminate the possibility that increased p16 expression was due to infiltration by p16-positive immune cells^37^, we isolated the stromal vascular fraction of the VAT and examined cells using CD45, an antigen associated with immune cells (Fig. 3i). The immune cell fraction had no difference in p16 expression, whereas non-immune cells from SD mice had increased p16 expression (Fig. 3j).

## Conclusion

Sleep disruption is endemic to modern society and is caused by many social circumstances such as shift work, newborn care, and chronic stress. Our study establishes a novel connection between a social determinant of health and aging. This data lays the groundwork to study the impact of senescent cells in those with frequently disturbed sleep. For example, chronic inflammation is exacerbated by the accumulation of senescent cells and is a key risk factor for many age-related diseases, such as heart disease^38,39^. Moreover, type 2 diabetes in humans has been linked to senescent adipose cells^40^. We uncover a potential target that could mitigate the systemic impact of sleep loss, as removing senescent cells can improve endothelial repair and metabolic dysfunction^41,42^. Furthermore, the incidence of sleep disturbances rises with advancing age in humans and this could be a driver for their increase senescent cell burden^7^. This exploration not only offers insights into the biological underpinnings of sleep but also holds promise for improving public health outcomes.

## Methods

### Mice

All animal experiments were conducted in accordance with the policies of the NIH Guide for the Care and Use of Laboratory Animals. Specific protocols used in this study were approved by the Institutional Animal Care and Use Committee of the Buck Institute.

Male C57BL/6J mice were purchased from the Jackson Laboratory (Bar Harbor, ME, USA). After arrival, mice were allowed to acclimatize for at least 7 days and remained in the same groups of four or five that they arrived in throughout the experiment.

### Sleep Disruption

Mouse weights were recorded before placing them in sleep-disruption (SD) or control cages, ensuring an equal starting weight average in each group. Mice were allowed to acclimate to new cages for at least 48 hours before sleep disruption began.

On the first day of sleep disruption, the machine’s spin speed was set to 3.5, corresponding to a 30-second bar rotation with random changes in rotational direction and an automatic 2-second speed difference enabled to delay acclimation. The spin speed was increased by 0.5 weekly due to reduced responsiveness, which is equivalent to about a 4-second increase in rotational speed (Supplementary Table 1). The bar rotation continued 24 h/day throughout the experiment, and mice were sacrificed from Zeitgeber time 3:30–5:15 by CO_2_, with control and SD mice in alternation to avoid circadian effects.

