## Supplementary figures and images for "Chronically disrupted sleep induces senescence in the visceral adipose tissue of C57BL/6 mice"

### Supplemental Figure 1

**Supplementary Figure 1**

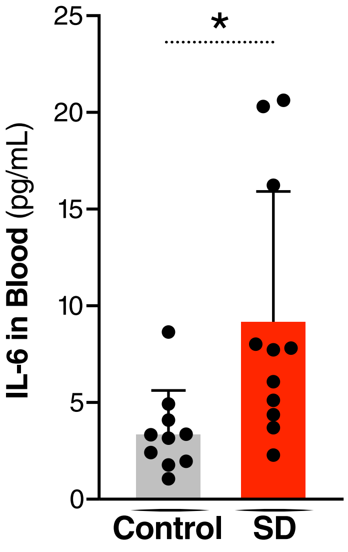


*Timonina 23.13E*

### Supplemental Figure 2

**Supplementary Figure 2**

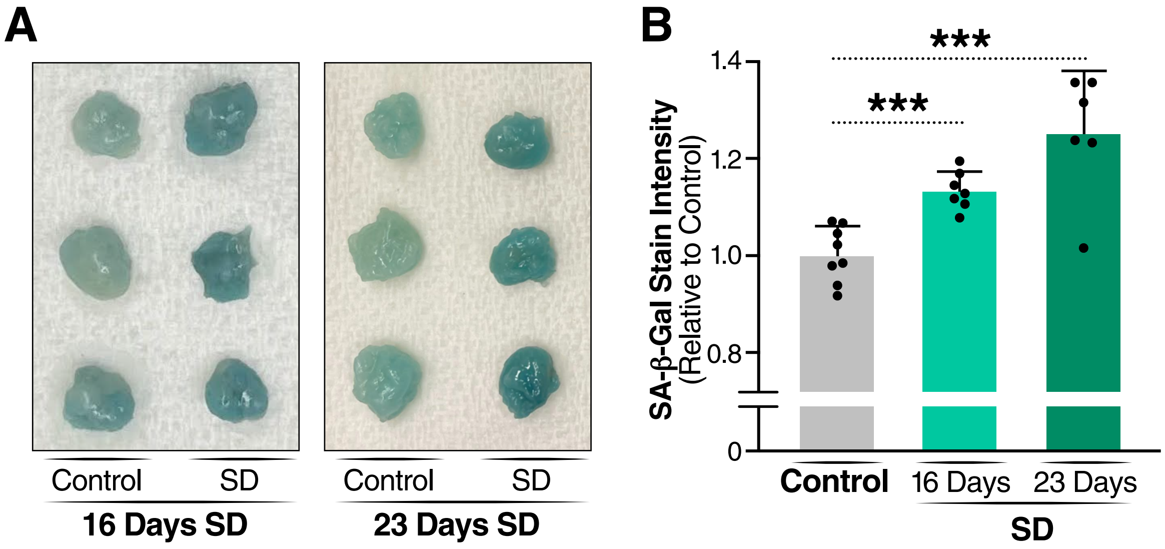


*Timonina 23.13F*
