## Supplemental Figure 2 Legend for "Chronically disrupted sleep induces senescence in the visceral adipose tissue of C57BL/6 mice"

Supplementary Fig 2.
Increased SA-β-gal activity precedes most changes in senescence gene expression. VAT tissue stained with X-gal (Cell Signaling) with representative images. Quantification of pixel intensity normalized to control average for each timepoint.[which stats test]
