## Supplemental Table 1 for "Chronically disrupted sleep induces senescence in the visceral adipose tissue of C57BL/6 mice"

### Supplementary Table 1

| SD Day | Speed Setting | Time to Complete One Rotation **(sec)** | |
|---|---|---|---|
| | | Fastest | Slowest |
| **1** | **3.5** | **27** | **32** |
| **7** | **4.0** | **20** | **28** |
| **14** | **4.5** | **17** | **24** |
| **21** | **5.0** | **15** | **21** |
| **28** | **5.5** | **11** | **16** |
