## Supplemental Table 2 for "Chronically disrupted sleep induces senescence in the visceral adipose tissue of C57BL/6 mice"

**Supplementary Table 2**

| Gene | Forward or Reverse | Primer Sequence (5′–3′) |
|---|---|---|
| **p16** | F | CCGTGTGCATGACGTGCGG |
| | R | AGTGGGGTCCTCGCAGTTCG |
| **p21** Variant 2 | F | AGGACGTCCCACTTTGCCAG |
| | R | CGGGACCGAAGAGACAACGG |
| **IL-1β** | F | AGCAGCTATGGCAACTGTTCCTG |
| | R | AGGACAGCCCAGGTCAAAGGT |
| **MMP3** | F | TGGGCCTGGAACAGTCTTGGC |
| | R | AGCAGCAACCAGGAATAGGTTGG |
| **ICAM-1** | F | GAACTGTGGCACCGTGCAGT |
| | R | GCAGGGTGAGGTCCTTGCCTA |
| **IL-6** | F | CCACTTCACAAGTCGGAGGCT |
| | R | TCTGCAAGTGCATCATCGTTGTTC |
| **TNF** | F | GCCTCTTCTCATTCCTGCTTG |
| | R | CTGATGAGAGGGAGGCCATT |
| **FBL** | F | CCGCAGGTTGCCTGAATCGC |
| | R | AAAGCCGCCTCTGCCACCAA |
| **SRP72** | F | AGAGTGCCACCCAGCAGACAG |
| | R | TCGTAGCGTTCCAGGCGGTA |
| **KPNA1** | F | CCTCTTCGTAGTGGCACCGGG |
| | R | TCCCTCCTCCTGCGCATCTCA |
| **GDF15** | F | GGTCGCTTCCAGGACCTGCT |
| | R | TGGCCGTGGGACCCCAATC |
| **SOD2** | F | GCCTGCTCTAATCAGGACCC |
| | R | GTAGTAAGCGTGCTCCCACA |
| **GSTP1** | F | ACGCAGCACTGAATCCGCAC |
| | R | CCTCACACCGCCCTCGAACT |
| **CAT** | F | TGTCGGACAGTCGGGACCCA |
| | R | GCCTCCGGTGGTCAGGACAT |
| HOUSEKEEPING GENES | | |
| **Actin** | F | ACTGTCGAGTCGCGTCCA |
| | R | GCGCAGCGATATCGTCATCCA |
| **HPRT** | F | CCCTGGTTAAGCAGTACAGCCCC |
| | R | GGCCTGTATCCAACACTTCGAGAGG |
